# Investigating social communication in mice: a Two-intruders test approach

**DOI:** 10.1101/2023.11.27.568787

**Authors:** Maryana V. Morozova, Lidiya V. Boldyreva, Maria A. Borisova, Elena N. Kozhevnikova

**Affiliations:** Scientific-Research Institute of Neurosciences and Medicine, Novosibirsk, Russia; The Federal Research Center Institute of Cytology and Genetics of the Siberian Branch of the Russian Academy of Sciences, Novosibirsk, Russia; Novosibirsk State Agrarian University, Novosibirsk, Russia

**Keywords:** social behavior, mouse model, mating, aggression, odor preference, testosterone

## Abstract

Understanding the complex dynamics of social communication behaviors, such as exploration, communication, courtship, mating, and aggression in animal models is crucial to reveal key neural and hormonal mechanisms underlying these behaviors. The Two-intruders test is designed to investigate residents’ behavior toward a male and female intruders. During this test imitating natural conditions, several aspects of social interaction were investigated: exploration, courtship, mating, and aggressive behavior. As mating and aggression involve overlapping neural circuits, the behavioral setup testing both behaviors is best at reflecting their competitive nature. Our findings demonstrate that male mice exhibit strong preference to communicate with a female intruder, which correlates with baseline testosterone levels of test males. Relevant female preference in the Two-intruder test was also found in BALB/c males. Behavioral breakdown revealed the anogenital sniffing as a key behavioral feature that discriminates test male behavior toward intruders of different genders. At the same time, female preference was accompanied by neuronal activation in the ventromedial hypothalamus. We demonstrate that odor recognition underlies preference toward females in male residents, as experimental anosmia reduced communication with a female intruder. However, there was no correlation between female animal preference in the contact Two-intruder test and smell preference in the social odor preference test. We assume the Two-intruders test setup to be a useful tool to study the neurological basis of social communication in animal models. Combined with odor preference tests, this experimental paradigm can help to decipher neural circuits involved in social deficiency phenotypes in animal models of human diseases.

**Significance Statement:** The Two-intruders test proves to be a highly reproducible and robust approach to assess social communication in mice. We demonstrate that the results obtained in this experimental setting replicate in different mouse groups and strains. This test is indispensable in studies assessing the competitive nature of male- and female-driven behaviors and the underlying neural mechanisms. Resident’s social interactions in the described here set up reflects odor processing and circulating testosterone – the key physiological drivers of animal communication with conspecifics. While easy to perform, this test provides a broad spectrum of behavioral patterns to study in the models of complex neurological diseases. Largely overseen in the literature as compared to the resident-intruder setup, the Two-intruders test can provide superior performance when used to understand the neural mechanisms of mating, aggression and social communication.

## Introduction

Social behavior is an integrative concept that involves multiple types of behavior and neural mechanisms behind it. Numerous human diseases are known to be associated with impaired social behavior including autism, anxiety, depression, dementia and other mental disorders (Baddeley, 2013; Frye, 2018; Keifer, Mikami, Morris, Libsack, & Lerner, 2020; Mendez et al., 2014; Tone, 2019). Basic aspects of human behavior are commonly studied in laboratory animal models, such as aggression, communication with live objects, maternal and defensive behavior, mating, and others (Fields, 2019; Lattal, 2001; Sare, Lemons, & Smith, 2021; Shemesh & Chen, 2023). It has been accepted that social interest, social recognition, and aggression are well suited to be tested in mice in the laboratory setting (Clipperton-Allen, 2023; Sare et al., 2021). These behaviors highly depend on perception of odor cues, as olfaction is a crucial sensory modality for rodents mediating social behaviors (Arakawa, Cruz, & Deak, 2011; J. Bakker, Leinders-Zufall, T., Chamero, P., 2022).

Major characteristics of social behavior are interactions between same- and opposite-sex animals. In a mouse brain, key structures that control interactions with male and female animals are well described. Social communication in mice begins with odor sensing in accessory olfactory system (AOS) and the main olfactory system (MOS) (J. Bakker, Leinders-Zufall, T., Chamero, P., 2022; K. Hashikawa, Hashikawa, Falkner, & Lin, 2016; Mohrhardt, Nagel, Fleck, Ben-Shaul, & Spehr, 2018). Most likely, MOS is responsible for the initial approach as it detects volatile odorants, whereas AOS detects urinary proteins and is involved in close contact (Arakawa, Arakawa, Blanchard, & Blanchard, 2007; Chamero et al., 2007; Luo, Fee, & Katz, 2003). Odor information from the main olfactory bulb is relayed to multiple downstream areas, including the cortical amygdala (СОА), which mediates approach and avoidance behaviors for innately attractive or aversive odors and retains the topographic organization of the olfactory bulb (Lin, Zhang, Block, & Katz, 2005). Mitral cells of the main olfactory bulb (MOB) project to several higher centers including the piriform cortex and the COA, which is further connected to medial and central amygdala, bed nucleus of stria terminalis (BNST), nucleus accumbens, and other limbic and infralimbic areas (J. Bakker, Leinders-Zufall, T., Chamero, P., 2022; Stamatakis et al., 2014; Yang, Karigo, & Anderson, 2022; Zhu, Tao, Huang, Qu, & Wang, 2023). Importantly, COA neurons express both: androgen and estrogen receptors suggesting that the COA cells’ response to odors may be under the control of the sex hormones (Y. Kim et al., 2015; Nodari et al., 2008; Yang et al., 2022). AOB projections are found to the medial amygdala, COA, bed nucleus of the accessory olfactory tract, and BNST (Mohedano-Moriano et al., 2007; Mohrhardt et al., 2018). BNST serves as a connection point that obtains input related to sex-specific olfactory signals and then projects them to and hypothalamic areas responsible for aggression and mating (K. Hashikawa et al., 2016; Lo et al., 2019; Yang et al., 2022). The medial preoptic area (MPOA) and ventromedial hypothalamus, ventrolateral subdivision (VMHvl), were shown to control mating and aggression programs in mice (Y. Hashikawa, Hashikawa, Falkner, & Lin, 2017; Karigo et al., 2021; Nair et al., 2023; Wei et al., 2018). VMHvl neurons are responsible for aggression and attacking and inhibit mating behavior via MPOA, whereas MPOA controls close investigation and mounting, while inhibiting aggression thought VMHvl (Y. Hashikawa et al., 2017; Wei et al., 2018). Strong stimulation of oestrogen receptor 1 (Esr1)-positive neurons within VNHvl evoked attack in male mice, and weak stimulation of these neurons resulted in mounting towards both males and females, as well as sniffing and close investigation, rather than attack (H. Lee et al., 2014).

To assess male-female and male-male behavior, a number of experimental paradigms were described (Clipperton-Allen, 2023; Kaidanovich-Beilin, Lipina, Vukobradovic, Roder, & Woodgett, 2011; D. G. Kim et al., 2019; Koolhaas et al., 2013; Tikhonova M.A., 2015). The initial preference of social odors is assessed in smell preference tests that involve an odor source or a choice of odors in the form of urine marks or soiled bedding samples. In this behavioral setting, a test animal is evaluated on its ability to memorize, discriminate and choose odors in terms of its relevance to the trait under study. For instance, male mice usually prefer female scent to male scents while estrous females’ odors are preferred to diestrous ones’; female mice prefer males of the same species, etc (J. Bakker, Leinders-Zufall, T., Chamero, P., 2022; Gheusi, 2008; Zolotykh M.A., 2017; Zou, Wang, Pan, Lu, & Xia, 2015). To understand further the nature of male-to-male and male-to-female social communication, contact tests are performed in home or foreign environment, which allows testing different aspects of social interactions and hierarchy (Blanchard, Wall, & Blanchard, 2003; D. G. Kim et al., 2019; Premoli et al., 2019). These are resident-intruder tests, involving interactions with same- or opposite-sex intruders (Koolhaas et al., 2013; Thomas, 1973). The resident-intruder test allows accession of natural behavioral expression of laboratory rodents in a semi natural laboratory setting. By recording the frequencies, durations, latencies and temporal and sequential patterns of all the observed behavioral acts, a detailed quantitative picture resident and also intruder behaviors can be obtained. Thus, a number of studies describe interaction with a male and a female in a resident-intruder paradigm in mice in different settings (Belousova I.I., 2009; Kuznetsova E.G., 2006; Leypold et al., 2002; Liu et al., 2011; Morozova et al., 2022; Norlin, Gussing, & Berghard, 2003; Zinck & Lima, 2013). However, given the competitive and mutually repressive nature of male- and female-derived cues in MPOA and VMH, simultaneous interaction with animals of both sexes represents social communication as it occurs in a natural environment.

Here we describe an experimental setup to perform a contact test with a male and a female intruder using male mice as residents in home environment. Our data suggest that this approach provides reliable and reproducible results in evaluating social interactions like investigation, preference, aggression, and mating while minimizing the effect of foreign environment.

## Methods and Materials

### 2.1 Laboratory animals and housing

The experiments were performed in the Scientific Research Institute of Neurosciences and Medicine. All procedures were carried out in accordance with Russian standards of Good Laboratory Practice (directive # 267 dated 19.06.2003 of the Ministry of Health of the Russian Federation) guidelines and the European Convention for the Protection of Vertebrate Animals used for Experimental and other Scientific Purposes. All procedures were approved by the Bioethical committee of SRINM (protocols #8 dated 15.08.2019 and #3 dated 19.05.2022).

The study utilized C57BL/6JNskrc (our in-house C57BL/6J sub-colony) and BALB/cNskrc (our in-house BALB/c sub-colony) mouse strains.

Adult 8–14 week-old mice were housed in groups of the same-sex siblings in open cages in 12 h/12 h light/dark photoperiod under standard conditions of a conventional animal house. Food and water were provided *ad libitum*. All animals were tested for pathogens according to Federation of European laboratory animal science association’s (FELASA) recommendations (FELASA et al., 2014).

### 2.2 Sexually experienced test males

To provide sexual experience, C57Bl/6JNskrc male mice at the age of 10-12 weeks were placed in individually ventilated cages (one male per cage) and co-housed with a sexually naïve oestrous female for 23±2 days. It is believed that the presence a female or sexual experience contributes to stronger territorial superiority and sexual motivation in males (Albert, Dyson, Walsh, & Petrovic, 1988; Amstislavskaya, Bulygina, Tikhonova, & Maslova, 2013). Then, the males were placed in fresh individually ventilated cages and single-housed for 7 days before the Two-intruders test.

### 2.3 Experimental groups

*Test C57Bl/6JNskrc males* (hereafter C57BL/6 males): To assess the reproducibility of the test, four independently tested groups of sexually experienced males were used at different times (n=9, 10, 11, and 10).

Group 1 (n=9) was used in the odor preference test (OPT) and the Two-intruders test

Group 2 (n=10) was used in the odor preference test (OPT) and the Two-intruders test

Group 3 (n=11) was used in the odor preference test (OPT), the Two-intruders test (with behavioral breakdown), and testosterone measurement.

Group 4 (n=10) was used in the Two-intruders test

An independent group of sexually experienced males (n=20) was tested in the Two-intruders test after anosmia induction.

An independent group of sexually experienced males (n=5 per group) was used for brain immunohistochemistry (IHC) analysis.

*Intruder BALB/cNskrc animals* (hereafter BALB/c): There were four independently prepared groups of 10 week-old animals used as intruders: n=12-13 males, and n=10-11 females for each Two-intruder test assay.

*Test BALB/cNskrc males* (hereafter BALB/c males): The one group of sexually experienced males, n=10 was tested in the Two-intruders test.

*Intruder C57Bl/6JNskrc animals* (hereafter C57BL/6): The group of 10 week-old animals was used as intruders in the Two-intruders test: n=12-13 males, and n=10-11 females.

### 2.4. The odor preference test (OPT)

To assess the smell preference of male mice toward male or female odor, a social odor preference test was used. The test was performed as follows: BALB/c female and male bedding samples were placed in two separate tea infusers (Ikea, art. #469.568.00) and introduced into the test animal’s cage. The animal was allowed to investigate the samples for 5 min. Characteristic “sniffing” (nose and whisker movements) were scored. The outcome is given as (time sniffing female or male bedding)/(total sniffing time) and expressed in percentages.

### 2.5. The Two-intruders test procedure

1. Place sexually experienced males into fresh cages at least 4 days prior to the test day. The two-intruders test in carried out in soiled cages, so bedding should not be changed at least 4 days before the test day.

*Note: Two-intruders test can also be performed in naïve males, if this is consistent with the objectives of the study. However, sexual and aggressive behaviors are less pronounced in naïve males*.

2. Prepare intruder animals one day prior to the test by marking their fur at the scruff or a tail with odorless food coloring (DyeCraft® Blue Food Coloring (USA), or similar) or cutting some fur on the back of dark-haired intruders to discriminate between males and females (only intruders of one gender are labelled)(Burn, Mazlan, Chancellor, & Wells, 2021). We suggest using intruder animal strain of the different fur color to distinguish animals during experiment. In the present study, C57BL/6 males were used as residents, whereas BALB/c males and females served as intruders and vice versa. Importantly, highly aggressive or extremely anxious strains should be avoided, as well as bigger strains. In our hands, both C57BL/6 and BALB/c animals appeared appropriate, even though BALB/c are known to be more anxious in other tests like open field test (Michalikova, van Rensburg, Chazot, & Ennaceur, 2010).

*Note: it is preferred choosing naïve males as intruders no older than 10-12 weeks old so that they are of comparable size with the residents or smaller and are not experienced to dominate or fight in a foreign environment. In case an intruder attacks a resident more than 5 times during the test, we suggest to interrupt the experiment and repeat with another intruder on the next day. Thus, preparing spare intruder males is recommended*.

3. The Two-intruders test is performed in home cage in the first dark hours of the diurnal cycle. This period seems optimal, as animals are most active in this phase. We suggest using red light since white light might induce anxiety and reduce social activity of test mice.
4. On the test day, remove cage covers, water bottles, food baskets, nesting and enrichment materials (if used) and install cages with test animals in the view field of the upright camera and the researcher. Make sure the video quality is high enough to allow further analysis of the recordings. Prepare timers and transparent plastic lid or equivalent material to cover the cages during testing.
5. Place a female and a male intruder into the test cage, cover with the transparent lid, start the camera and the timers, let the animals interact for 15 minutes. After the test is finished, place intruders into fresh cages to discriminate them from unused ones. Return water, food and nesting material back to the residents’ cage. Wipe the transparent thermoplastic lid with 70% ethanol and let it dry while preparing the next animal.
6. Analyze behavior during the test visually or later using video recording, or behavior recognition software. In this work we calculated chasing, nasal sniffing, anogenital sniffing, grooming, mating (mounting with pelvic thrusting movements), attacks and fighting visually. Two independent researchers performed the calculations, and their results matched. The data were not blinded in these experiments, as there were no differences between the test groups.

### 2.6. Intranasal administration of zinc sulfate

To induce anosmia, C57BL/6 mice were nasally treated with 10% ZnSO4 (AppliChem GmbH, Germany) in PBS in a volume of 50 µl one day prior to testing behavior as recommended (McBride, Slotnick, & Margolis, 2003). Control animals were nasally treated with the equal amount of PBS.

### 2.7. Immunohistochemistry

Neuronal activation was assessed using IHC staining for neuronal cell activation marker protein c-Fos according to Lee et al. (H. Lee et al., 2014) with the following modifications. Adult male mice were single-housed prior to the neuronal activation in their home cages. A male intruder or a female intruder or a tissue with the neutral odor (lemon) was introduced as a stimulant for 10 minutes. After 1 hour mice were anesthetized intraperitoneally as follows: Domitor (75 µL/10 g weight; Orion Pharma, Espoo, Finland) dissolved at 0.05 mg/ml in PBS was injected at 10 µl/g; 15 min later Zoletil (60 µL/10 g weight; Virbac Sante Animale, Carros, France) in concentration of 5 mg/ml was injected at 10 µl/g. Intracardiac perfusion was performed using 15 ml of PBS and 15 ml of 4% PFA (AppliChem GmbH, Germany) per animal as described earlier (Glinskikh et al., 2021). Brains were post-fixed overnight in 4% PFA, and then kept in 15% sucrose in PBS solution for 24 h and in 30% sucrose in PBS solution for another 24 h at 4⁰C. Then brain were embedded in Tissue-Tek® O.C.T. Compound (Sakura Finetek, USA) and frozen in liquid nitrogen. 30 μm sections of brain in the ventromedial nucleus of the hypothalamus (VMH) region were prepared using 550 HM Microm cryostat (ThermoFisher Scientific, USA). Sections were incubated in 1% bovine serum albumin (BSA) + 0,15% Triton X-100 in PBS for 2 h. Sections were washed three times for 5 min in PBS + 0.1% Triton X-100 (PBST). Primary antibodies to c-Fos (#sc-166940, Santa Cruz, USA), in 1:300 dilution, were incubated overnight at 4°C in PBST+0,1% BSA and washed three times for 5 min with PBST. Secondary antibodies (# R37116 Anti-Rabbit IgG (H+L) Cross-Adsorbed Secondary Antibody, Invitrogen, USA), in 1:500 dilution, were incubated for 2 h at room temperature in PBST + 0,1% BSA and washed three times for 5 min with PBST. Then brain sections were incubated in PBS with 0,15 µg/ml DAPI for 1 h and washed with PBS once. Stained sections were stored in Vectashield® Antifade (Vector Laboratories, USA) on glass slides at 4°C. Images were obtained using ZEISS AxioScope.A1 with oil immersion 40×/1,3 and 100×/1,4 plan-apo lenses and the ZEN 2012 software. The number of activated neurons was counted on brain sections in the ventromedial nucleus of the hypothalamus (VMH), the VMH nuclei were mapped according to the Allen Brain Atlas. Quantification was performed using ImageJ software. At least 30 independent measurements were made for 5 animals per group. Average number of activated neurons was calculated for Control (Lemon odor) group in the VMH region. The number of activated cells for male intruder or female intruder groups was calculated as follows: Number of c-Fos^+^ cells — Average number of activated cells in the Control group.

### 2.8. Testosterone measurement

The level of testosterone in blood serum was measured by ELISA using a commercial testosterone-ELISA-BEST kit according to the manufacturer’s protocol (X-3972, VectorBest, Russia). A Berthold TriStar LB-941 multimode reader was used foe detection (Berthold Technologies GmbH & Co., Germany).

### 2.9. Statistics

The data are presented in boxplots showing the median, 25th and 75th percentiles, minimum, maximum, and outliers. Individual animals’ measurements are represented as data points on the graph. The data were tested for normality using the Kolmogorov–Smirnov test. Normally distributed data were analyzed by Student’s t-test test for dependent and independent samples. Data involving multiple factors were analyzed using analysis of variance (ANOVA) followed by Tukey HSD test. For not normally distributed data, Mann-Whitney U test was used for independent samples and Wilcoxon Matched Pairs test was used for dependent samples. Categorical data was analyzed using Fisher exact test. Pearson correlation analysis was used to assess the correlation. Principal component analysis (PCA) was performed in STATISTICA.12 software (StatSoft TIBCO Software), data significance in principal components (PC) was evaluated using Student’s t-test for independent samples. The value of p < 0.05 was considered significant.

## Results

### 3.1. Male mice prefer interacting with female intruders in the Two-intruders test

We evaluated the test male social interactions in the Two-intruders test using C57BL/6 mice and measured time of communication with a female or a male intruder in seconds and in percentages of total social interaction. We also counted aggression, mounting and calculated the number of contacts with a male and a female intruder. The data shown in Fig. 1A-D shows joint results from four independent experiments. The number of contacts was assessed in two independent experiments and Fig. 1E demonstrates the combined outcome of the two. We found that male resident mice preferred to interact with a female intruder. Test males spent more time with a female intruder as measured in absolute values (n=40, *t*=-11.5, *p*<0.001, Student’s t-test for dependent samples, Fig. 1A) and as a percentage of total social interaction (n=40, *t*=-12.18, *p*<0.001, Student’s t-test for dependent samples, Fig. 1B). Aggression was evaluated as the number of aggressive acts (fighting and biting) toward a female or a male intruder, and resident mice were more aggressive to a male intruder (n=40, *p*<0.05, Fisher exact test, Fig. 1C). Likewise, sexual behavior was assessed as the number of mating acts (mountings with pelvic thrusting movements), and test males normally can attempt to mount a male, but prefer to mount a female (n=40, *p*<0.001, Fisher exact test, Fig. 1D). The number of contacts also supports the overall interest to female intrudes, as test males prefer to communicate with females more often (n=20, *t*=5.33, *p*<0.001, Student’s t-test for dependent samples, Fig. 1E).

**Figure 1.**
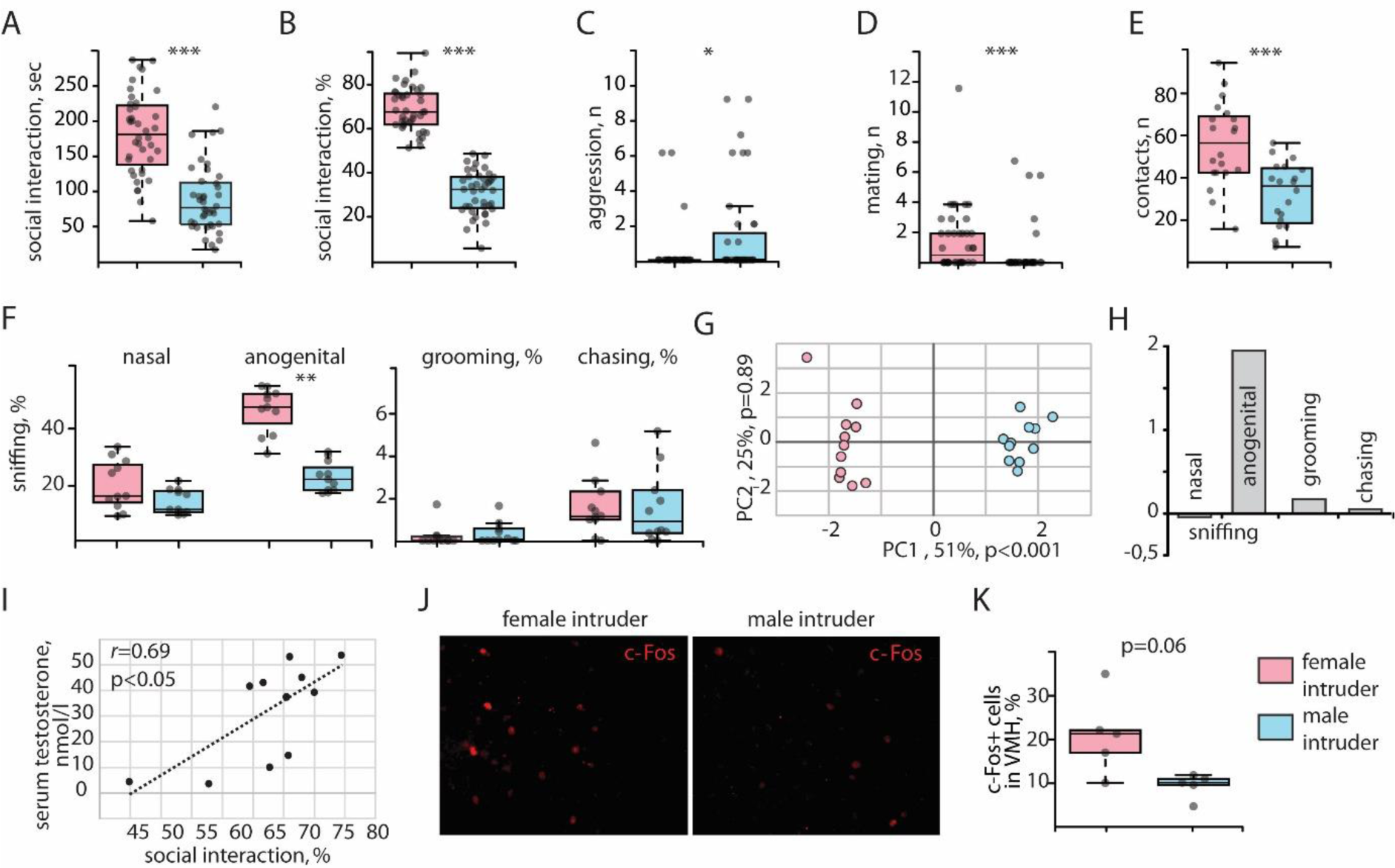
Test male interaction with male and female intruders in Two-intruders test analysis. **А**. Relative duration of social interaction with intruders, % of the total interaction time, n=40; **В**. Duration of social interaction with intruders, seconds, n=40; **С**. Count of attacks by test male on intruders, n=40, **D**. Count of mounting episodes on intruders by test male, n=40; **E**. Count social interaction episodes with intruders by test male, n=20; **F**. Breakdown of test male social interaction: relative duration of naso-nasal sniffing, anogenital sniffing, grooming and chasing of intruders, % of the total interaction time, n=11; **G.** PCA analysis of behavioral traits, n=11. **H**. Score contribution to principal components. **I**. Correlation curve between the level of testosterone in the test male serum and the relative duration of female sniffing, n=11, **J**. IHC staining of in VMH sections for c-Fos after presentation of a female and a male intruder to the test male, n=5*30 cells; **K.** Count of the c-Fos-positive neurons in VMH sections of test male brain after presentation of a female and a male intruder to the test male, n=5. * = p < 0.05, ** = p < 0.01, *** = p < 0.001, if not indicated otherwise.

Next, we investigated the spectrum of other social behaviors found in the Two-intruders test in one of the described above experiments. The breakdown of communicational patterns revealed that anogenital and nasal sniffing are prevalent social activities during the test. Minor activities were chasing and grooming of intruders. Strong female preference was only found for the anogenital sniffing (n=11, *Z*=2.93, *p*<0.01, Wilcoxon Matched Pairs test, Fig. 1F). PCA analysis of the behavioral breakdown revealed that social communication patterns toward female and male intruders differ significantly in the first principal component (n=11, *t*=-27.99, *p*<0.001, Student’s t-test for independent samples, Fig. 1G). However, the main factor positively contributing to this difference was the anogenital sniffing (Fig. 1H), revealing this parameter to be key in behavioral manifestation of social preference.

We also evaluated how accurately the Two-intruders test reflects the physiological state of the test male in terms of circulating testosterone - the major driver of mating and aggression behavior. Thus, we performed correlation analysis of testosterone with female preference and found that baseline plasma testosterone in C57BL/6 males strongly correlates with total duration of time spend with the female in the Two-intruders test (r=0.69, p<0.05, Spearman’s correlation coefficient, Fig. 1I).

Social interactions including mating and aggression are translated through the neuronal activation of the deeper brain structures, and particularity the hypothalamic nuclei. To evaluate the extent of the observed social preference in resident, we detected activation of the VMH via c-Fos immunohistochemistry after direct contact with either a male or a female (Fig. 1J). The number of c-Fos-positive cells in the VMH in test males tended to be higher upon communication with a female intruder (p=0.06, Mann-Whitney U test, Fig. 1K).

### 3.2. Female bias relies on odor discrimination, but does not correlate with the odor preference itself

The olfactory cues are known to guide female preference in mice under natural conditions. Thus, we induced anosmia in the test males by intranasal instillation of zinc sulfate to reduce odor discrimination (McBride et al., 2003). As result, male mice preferred female intruders at a lesser extent after the reduction of odor perception (Fig. 2A). ANOVA revealed the influence of the “intruder gender” factor (n=20, F(1,34) = 301,34, p<0.001) and the interaction of the “intruder gender” and “ZnSO_4_” factors (n=20, F(1, 34) = 150,99, p<0.001). Post Hoc analysis revealed that even though ZnSO_4_-treated males still preferred females (p<0.01, Tukey HSD test), they did so at a significantly lower rate than their untreated counterparts (p<0.001, Tukey HSD test). We then questioned whether social odor preference test could fully substitute for the Two-intruders test in terms of female preference by male intruders. In order to test this hypothesis, we compared the results of these two tests run consequently using the same untreated animals and discovered that bias toward a female intruder generally does not correlate with female smell preference in a given male (n=30, *r*=0.12, p=0.47, Spearman’s correlation coefficient, Fig. 2B). Importantly, these same males significantly preferred female odor in the odor preference test (n=30, *t*=12.46, *p*<0.001, Student’s t-test for dependent samples Fig. 2C).

**Figure 2.**
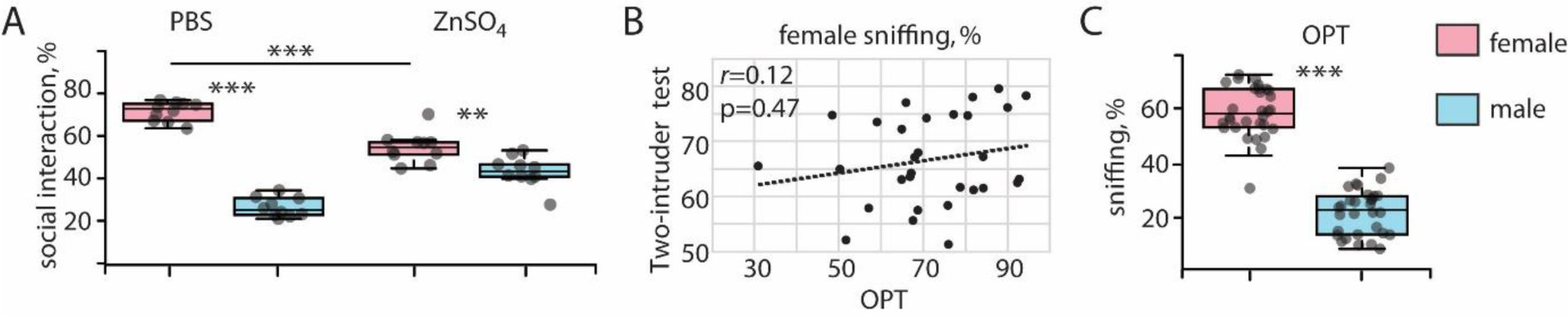
Olfaction role in the Two-intruder test. **A.** Relative duration of test male social interaction with female and male intruders after intranasal instillation of zinc sulfate, % of the total sniffing time, n=10 per group. ** = p < 0.01, *** = p < 0.001. **B**. Correlation curve between the relative duration of test male social interaction with female and the relative duration of female sniffing in odor preference tests (OPT), n=30; **C**. Relative duration of test male sniffing female and male odors in OPT, n=30.

### 3.3. The Two-intruders test provides robust and reproducible results

In Fig. 1 we demonstrated the combined results of 4 independent experiments using different groups of C57BL/6 test males. At the same time, an important characteristic of an experimental assay quality is reproducibility. To demonstrate the robust and reproducible results obtained in the Two-intruders test, we show that four independent replicas performed at different times with independent groups of animals agree on the results and show no statistically significant differences between the groups (Fig. 3A). ANOVA revealed the effect of the “intruder gender” factor (n=9-11 per group, F(1,72) = 328.29, p<0.001) and the interaction of the “intruder gender” and the “group” factors (n=9-11 per group, F(3,72) = 4.45, p<0.01) on female/male intruder preference. Post-hoc analysis revealed a strong preference for intruder females in all four test replicates (p<0.001, Tukey HSD test) and no differences between the groups. The same is true for the absolute time of interaction with either of the intruders (Fig. 3B). ANOVA revealed only the effect of the “intruder gender” factor (n=9-11 per group, F(1,72) = 492.83, p<0.001), and no effect of the group was found. Neither there was an effect of the “group” factor on total duration of social interaction in these replicates (Fig. 3C), however there was a noticeable dispersion of this parameter. Thus, the Two-intruders test provides reproducible results showing that resident males highly significantly prefer a female intruder for social communication.

**Figure 3.**
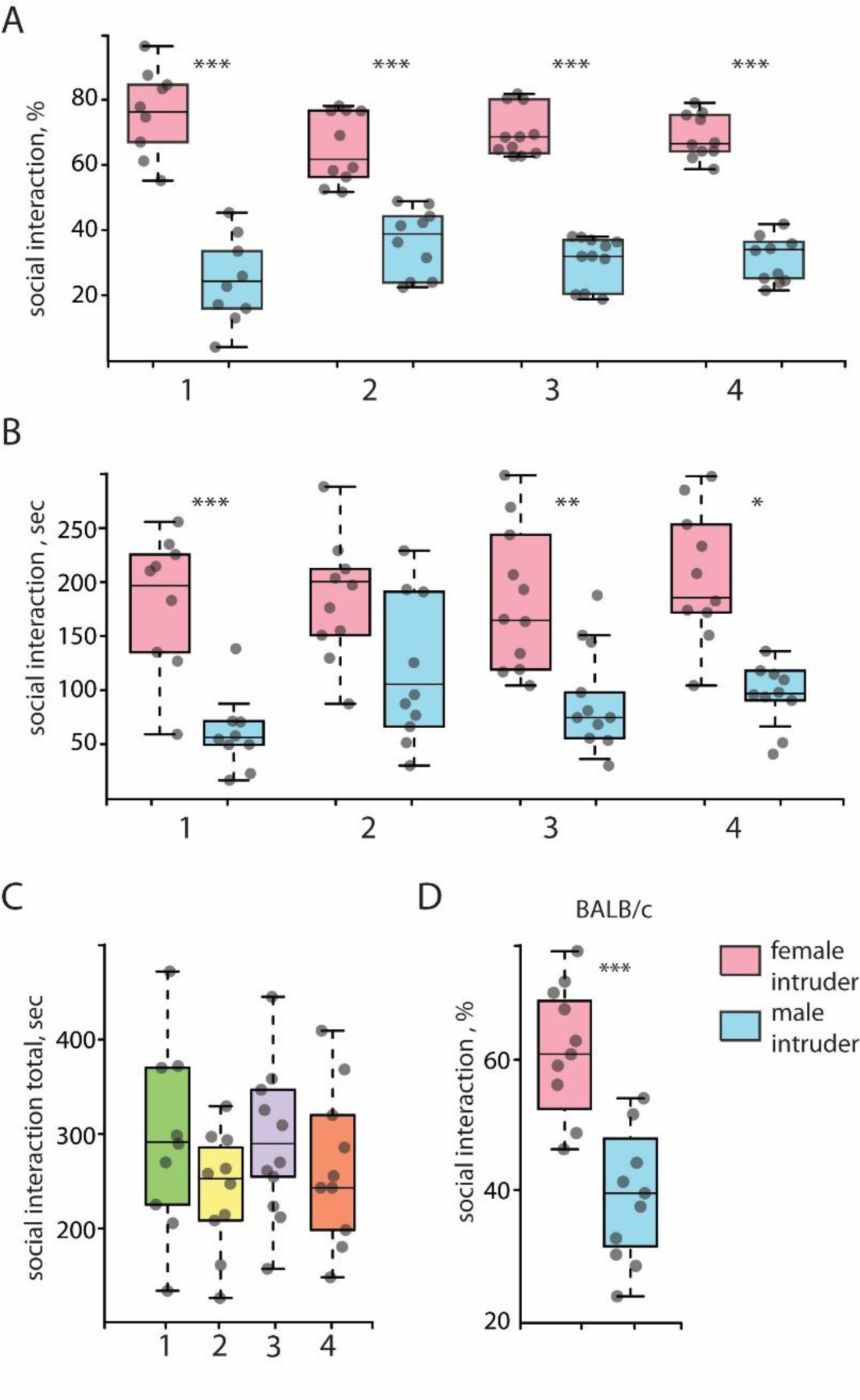
Two-intruders test reproducibility. **A.** Relative duration of test male social interaction with female and male intruders in four biological replicates, % of the total sniffing time, n=9-11; **B**. Duration of test male social interaction with female and male intruders in four biological replicates, n=9-11; **C**. Total duration of test male social interaction with intruders in four biological replicates, n=9-11. **D**. Relative duration test male social interaction with female and male intruders in BALB-c strain, % of the total sniffing time, n=10. ** = p < 0.01, *** = p < 0.001

Different mouse strains exhibit controversial behavioral characteristics in the some behavioral tests, reflecting the lack of their universal utility (Amstislavskaya & Khrapova, 2002). To explore further the robustness of the aforementioned experimental procedure, we applied it to another commonly used inbred laboratory mouse strain BALB/c. Our results demonstrate that BALB/c males also significantly prefer to interact with a female intruder rather than the male one (n=10, t=5.36, p<0.001, Student’s t-test for dependent samples, Fig. 3D).

## Discussion

The Two-intruders test reported in this study is based on the previously described mating choice assays and our own design published earlier (Amstislavskaya et al., 2013; Koolhaas et al., 2013; Kuznetsova E.G., 2006; Leypold et al., 2002; Liu et al., 2011; Morozova et al., 2022; Norlin et al., 2003). For instance, Leypold and colleagues used the Two-intruders test together with separate resident-intruder aggression and mating tests with either a male or a female intruder alone. They demonstrated the key role of VNO, and particularly, a TRP2 cation channel in odor processing and specialization of downstream behavioral patterns depending on the intruder gender (Leypold et al., 2002), the authors define this experimental paradigm as a “mating choice test” since it reflects increased sexual behavior toward male intruders in *trp2* mutants. Likewise, Liu and co-authors also refer to this setting as a “mating choice test” and use it do demonstrate sexual interest of serotonin-deficient mice indiscriminately to both male and female intruders (Liu et al., 2011). Later it was shown that mounting a male intruder does not necessarily reflects the intent and neural encoding of sexual behavior, and so far can only be discriminated by ultrasonic vocalizations that accompany mounting behavior (Karigo et al., 2021). For this reason, we suggest that the proposed experimental setup remains within the resident-intruder paradigm, as even wild-type mice exhibit some mounting activity toward male intruders, which hardly reflects sexual interest (Fig. 1D).

The setting for the Two-intruders test is relatively simple, and most researchers can perform it in similar conditions, with differences in the intruder choice and sexual maturity of the resident. It has been shown that experienced resident males exhibit pronounced aggression and mating as compared to the naïve residents (Amstislavskaya T.G., 2018; Karigo et al., 2021; Leypold et al., 2002), so that we suggest experienced males for the test as it provides wider dynamic range to compare social behavior between the control and test groups. Obviously, this recommendation does not apply to studies aiming social maturation as their objective or similar investigations.

When it comes to the male intruder choice, many experimenters tend to choose “comfortable” intruders like vasectomized males or sexually immature animals swabbed with a mature male’s urine (Fraser & Shah, 2014; D. L. Lee & Wilson, 2012). The reason to do so is to minimize the interference of a male intruder with the residents’ actions, which are mainly interrupted by the intruders’ aggression and attacks. However, a small immature male intruder might be a vulnerable target for an extremely aggressive resident ending up severely injured by the first third of the test. Thus, we avoid juvenile intruders in the described above setting. Adult vasectomized intruders appear a good alternative, but it implies a designated animal group to undergo surgical procedure for the purpose of this test only. To reduce labor, cost and animal misuse, we prefer intact naïve males as intruders, so that they can be used later in other experiments. Our results demonstrate that both BALB/c and C57BL/6 mice are suitable for the Two-intruders test as male intruders (Fig. 1,3), which agrees with the results obtained earlier (Morozova et al., 2022). Moreover, BALB/c intruders provoke efficient response in the resident-intruder assay (Amstislavskaya T.G., 2018; Amstislavskaya & Khrapova, 2002; Novikov S.N., 1988).

The Two-intruders test allows observing and measuring a number of behavioral parameters, which can be combined for a specific purpose of the study. It is mostly used to evaluate the test male’s ability to discriminate between males and females, so the characteristics of such discrimination are of the highest value. Apart from mating and aggression that clearly differentiate social behavior toward a male and a female (Fig. 1C-D), anogenital sniffing appears as another social activity that should be taken into account while evaluating social deficiency phenotypes in mice. The breakdown of behavioral pattern during the 15-minute Two-intruders test reveals that anogential sniffing of a female is the predominant activity of the resident male. It was the only additional parameter tested to contribute significantly into social discrimination between male and female intruders (Fig. 1G,H), which aligns well with currently accepted concept of mate choice in mice (Lenschow, Mendes, & Lima, 2022). Given mentioned above, we consider anogenital sniffing as a key measure of sexual preference.

Among most widely assessed parameters in resident-intruder tests are aggression and mating, both of which are under control of steroid hormones. It is known that estrogen-, androgen- and progesterone receptors play an important role in the control of reproductive behavior, and are widely distributed in the brain (J. Bakker, 2022; Balthazart, 2021; Kight & McCarthy, 2020). Here we show that blood testosterone levels in the test males strongly correlate with their preference towards a female intruder (Fig. 1I). This result is consistent with previous studies demonstrating the primary role of testosterone in female preference and sexual behavior (Amstislavskaya & Popova, 2004; David, Wyrosdic, & Park, 2022; Henley, Nunez, & Clemens, 2011; Pierman et al., 2008). Thus, the Two-intruders paradigm clearly reflects hormonal regulation behind mating behavior and can be utilized to assess the role of steroid metabolism and the associated deficits in social communication.

Mate preference can also be assessed in the indirect odor preference tests where male mice tend to prefer female’s odor to the male’s one and describe appetitive behaviors in social communication (Arakawa et al., 2011; Kaidanovich-Beilin et al., 2011; Zolotykh M.A., 2017). While the Two-intruder test largely reflects the consummatory phase of social interaction. We then questioned whether odor preference correlates with the intruder choice using the same group of animals. Although wild type C57BL/6 mice generally prefer female odor in the odor preference test and interacted more with females in the Two-intruders test, these two parameter did not correlate being assessed per individual animal (Fig. 2B). This result suggests that despite the fundamental role of odor recognition in social communication in mice (Fig. 2A), the neural encoding of odor processing and direct encounter are different in nature as has been reported (Bathellier, Buhl, Accolla, & Carleton, 2008; Saha et al., 2013; Smear, Shusterman, O’Connor, Bozza, & Rinberg, 2011). Our previous studies also demonstrate the essential difference in social preference in odor- and intruder-based tests. For instance, we earlier observed lack of female preference in the odor-based test with no changes in the Two-intruders test in chemically-induced model of chronic colitis (Borisova et al., 2020). At the same time, a transgenic model of chronic colitis exhibited full recognition of odors, but totally lacked female preference in the resident-intruder paradigm (Morozova et al., 2022). These behavioral differences finely reflected the diverse neurochemical patterns that accompanied chronic colitis in these two models. Therefore, a combination of these two tests appears as a powerful instrument to evaluate subtle changes within neural network that encodes social communication in the disease models.

## Acknowledgements

Fluorescent microscopy was performed at the Microscopy Center of the Institute of Cytology and Genetics, SB RAS, Russia (https://ckp.icgen.ru/ckpmabo/).

